# Delayed maturation of inner hair cell ribbon synapses in a mouse model of fragile X syndrome

**DOI:** 10.1101/2025.03.31.646200

**Authors:** M. Chojnacka, A. Skupien-Jaroszek, M. Dziembowska

## Abstract

Clinical features of FXS phenotype include intellectual disability, repetitive behaviors, social communication deficit and also commonly manifested auditory hypersensitivity for acoustic stimuli. The electrophysiological studies have shown that FXS patients and *Fmr1 KO* mice display improper processing of auditory information in the cortical areas of the brain and the spiral ganglion of cochlea. Synapses formed by spiral ganglion neurons on sensory hair cells (HC) are the first connection on the path that conveys the auditory information from the sensory cells to the brain. We confirmed the presence of fragile X messenger ribonucleoprotein (FMRP) in the inner hair cells of the cochlea. Next, we analyzed the morphology of IHC ribbon synapses in early stages of postnatal development (P5, P14) and detected their delayed developmental maturation in *Fmr1 KO* mice. Interestingly the ultrastructure of inner hair cell ribbon synapses studied by electron microscopy in the adult mice have shown no specific dysmorphologies. Delayed maturation of presynaptic ribbons of auditory hair cells in *Fmr1 KO* mice may contribute to abnormal development of circuits induced by auditory experience.

## Introduction

Fragile X syndrome (FXS) is a genetic disorder caused by transcriptional silencing of the FMR1 gene on chromosome X (Hagerman et al., 2017, Hagerman et al., 2005). The FMR1 gene contains nearly 50 CGG trinucleotide repeats in the 5’ untranslated region, but in FXS the repeats expand to over 200, this leads to hypermethylation in the expanded CGG repeats and no expression of FMRP protein (Crawford et al., 2001). FXS is the most common monogenic cause of inherited intellectual disability and autism spectrum disorder (Sinclair et al., 2017). FXS patients are often characterized by poor language development and prominent sensory abnormalities (Hagerman et al., 2017, Rais et al., 2018). In both FXS patients and animal models, studies show auditory hypersensitivity, impaired habituation to repeated sounds, reduced auditory attention, and difficulties in complex listening, phenotypes that may also be associated with problems in social interactions and language development (Rotschafer and Razak, 2013, Rotschafer and Razak, 2014, Rais et al., 2018, Razak et al., 2021).

The critical element of mammalian auditory system located in the organ of Corti are hair cells, the auditory receptors that form synapses with spiral ganglion neurons and transform mechanical vibrations into electrical signals (Wichmann and Moser, 2015). Two types of sensory hair cells are located in the organ of Corti, the inner hair cells IHC and outer hair cells OHC connected to SGNs by glutamatergic ribbon synapses, which are essential for faithful synaptic transmission (Glowatzki and Fuchs, 2002). The IHCs possess highly specialized presynaptic active zones (AZs) called synaptic ribbons with the characteristic RIBEYE protein stem and synaptic vesicles docked into it. Ribbon synapses of cochlear inner hair cells (IHCs) undergo molecular assembly and extensive functional and structural maturation before hearing onset. Recent studies show that aberrations in ribbon synapses are responsible for hearing loss (Liberman and Kujawa, 2017).

One of the hallmarks of neurodevelopmental disorders such as FXS is cell and circuit hyperexcitability (Goncalves et al., 2013, Zhang et al., 2014) often associated with such neurological symptoms as hypersensitivity, hyperarousal, hyperactivity, anxiety, and seizures (Penagarikano et al., 2007, Braat and Kooy, 2015) In Fmr1 knock-out mice, a model of FXS, abnormal dendritic spines with long, thin and immature morphology have been described (Comery et al., 1997). However, little is known about the morphology and neuronal connections between spiral ganglion neurons and inner hair cells. This led us to investigate the abundance of synaptic connections in hair cells in *Fmr1 KO* mice compared to wild-type mice.

In this study we performed the morphological analysis of IHCs in the wt and *Fmr1 KO* mice cochlea using complementary research methods that included confocal and electron microscopy imaging. This allowed us to study the morphology of the IHC in the organ of Corti and the fine structure of their ribbon synapses. We found no visible morphological differences in IHC morphology, distribution of mitochondria, the number of presynaptic active zones and postsynaptic neurites of SGNs when we compared *Fmr1 KO* and WT mice. The ultrastructure of reconstructed ribbons did not show any particular dysmorphologies or differences in the number of synaptic vesicles. However, using the immunofluorescent staining of presynaptic ribbons preformed on three developmental stages spanning the prehearing P5, hearing onset P14 and adults we identified a delayed formation of ribbon synapses of cochlear inner hair cells.

## Material and methods

### Animals

*Fmr1 KO* and wild type mice with background FVB strain (P5, P14 and adult) were used for experiments. Before the experiment, the animals were kept in the laboratory animal facility (Faculty of Biology, University of Warsaw) under 12-h light/dark cycle with food and water available *ad libitum*. The animals were treated in accordance with the ethical standards of European and Polish regulations.

### Immunofluorescence staining of hair cells of organ of Corti

To obtain inner ears, animals were sacrificed according to approved humane protocol and the inner ears were dissected from the skull. Round and oval windows were opened, and bone over the apical turn removed to allow rapid fixation of 4% paraformaldehyde with 4% saccharose in PBS through the cochlear for 5 min. Next, cochleas were transferred into PBS, to obtain their apical turn. Bone over the middle-ear-facing portion of the cochlear spiral and the tectorial membrane were removed by fine forceps. Following tissue were fixated in 4% paraformaldehyde with 4% saccharose for 40 min. For blocking and permeabilization, cochleas were incubated in PBS with 0,5% Triton X-100 and 5% BSA for 1 h, with gentel shaking. Next, tissue was incubated with primary antibodies at 4°C for overnight, and secondary antibodies Alexa Fluor (1:1000; Invitrogen) at room temperature for 2h. The following antibodies and fluorescent dyes were used: mouse IgG1 Anti-CtBP2 Clone 16/CtBP2 (1:500; BD Transduction Laboratories), mouse IgG2a anti-GluA2 (1:200; Millipore); rabbit anti-MyosinVI (KA-15) (1:1000, Sigma); mouse IgG2a anti-Bassoon (1:500, Santa Cruz); rabbit anti -FMRP (D14F4) (1:100, Cell Signaling); Phalloidin 405 nm (1:1000, Abcam), rabbit anti –Tom20 (FL-145) (1:250, Santa cruz). After immunostaining, cochleas were mounted on microscope slides in ProLong Gold antifade reagent (Invitrogen), and coverslipped.

### Confocal microscopy

Z-stacks from selected cochlear regions from each ear were obtained on confocal microscope (Zeiss Axio Imager Z2 LSM 700) with oil-immersion objective 63 x and 1× digital zoom. Images were obtained in a 2048 × 2048 frame (pixel size = 0.05 μm in x and y). Care was taken to minimize pixel saturation in each image stack. Each stack contained the entire synaptic pole of eight to eleven inner hair cells as viewed from the surface of the organ of Corti. The same microscope settings were used for all sections and all repetitions.

### Image analysis

The images were analyzed in the 3D Object Counter (https://imagej.net/3D_Objects_Counter) application running in the ImageJ/Fiji. Intensity threshold settings were adjusted to identify the vast majority of synaptic densities, while excluding non-synaptic staining. The histogram depicting the frequency of ribbons as a function of the volume of the presynaptic, anti-CtBP2 stained puncta was performed using GraphPad Prism version 7.00 for Windows, GraphPad Software, La Jolla California USA, www.graphpad.com.

### Electron microscopy

To maintain proper structure of hair cells in samples collected from adult mice, we removed the membrane from the round window and the stapes footplate of the oval window, and opened a small hole in the bone at the apex of the cochlea. Next, solution for fixation were gently perfused by syringe and perfusion-fixed on ice over-night.

### Transmission electron microscopy

The whole cochlea was cut out together with the bone capsule and transferred 2.5% glutaraldehyde (Electron Microscopy Sciences), 2% formaldehyde (fresh from paraformaldehyde (EMS) in PBS at 4 ° C overnight. After washing with PBS, the tissue was incubated in 1% osmium tetroxide and dehydrated in graded aqueous ethanol solutions from 25% to 96% (each for 10 minutes) followed by 100% ethanol (two lesions each for 10 minutes). Infiltrated with a mixture of 25% and epoxy resin 30 minutes on ice, 50% epoxy resin for 2 hours, 75% and 100% for 24 hours at room temperature Then, the samples in 100% epoxy resin were transferred to a flat embedding mold and placed in an oven at 60 ° C for 48 h. Samples were cut on an ultramicrotome (Leica) by diamond Diatome knife (Ted Pella) into 100 μm thick slices. The ribbon was transferred to tungsten coated copper meshes and incubated with 1% uranyl acetate followed by lead aspartate. Electron micrographs were obtained on a transmission electron microscope JEM 1400 (JEOL).

### Serial block face scanning electron microscopy (SBF-SEM)

Tissue preparation using this protocol allowed for partial reconstruction of hair cell ultrastructure (Bullen et al., 2015). After fixation (cold 2% paraformaldehyde, 2.5% glutaraldehyde solution in PBS) and decalcification cochleas were transferred to 0.1% tannic acid (EMS,USA) in PBS and incubated in this solution overnight at 4°C. Samples were processed for SBF-SEM following the method of Deerinck et al. (2010). Samples were trimmed and mounted into pins. Smooth surface of sample to SBF-SEM was obtained by using ultramicrotome (Leica) and diamond knife (Ted Pella). Next, the sample were painted with silver paint (Ted Pella) for enhance conductivity. Samples were imaged by SBF-SEM Sigma VP 3View at the Nencki Institute of Experimental Biology through the technical platform (Laboratory of Imaging Tissue Structure and Function). Scanning parameters: variable pressure 25Pa, EHT 3kV, aperture 30um, dwell time 2-5us and 7-9us. Blocks were cut at 50 nm and 150 nm, and the exposed block face imaged with a pixel size of 3,7 -4,2 nm and 10 – 11 nm for synapse and inner hair cells images, respectively.

### The analysis of hair cells ultrastructure

3D models for whole cell, cellular organelles such as mitochondria and ribbon synapses in the hair cells of WT and *Fmr1 KO* mice were prepared using Reconstruct software (Fiala, 2005) http://synapses.clm.utexas.edu/tools/reconstruct/reconstruct.stm.

### Statistics

All the statistical details including statistical tests used and number of biological replicates are described in the Figure Legends. The analyses were performed using Prism

10.4.0 software (GraphPad, San Diego, CA, USA).

## Results

### Expression of FMRP in the hair cells of the organ of Corti

We examined the expression of FMRP in the hair cells of mice. Staining was performed on tissue isolated from the cochlea of P5 and adult WT and *Fmr1 KO* mice (n = 2 per genotype for P5, n = 3 per genotype for adult mice). We used an anti-CtBP2 antibody as a marker for nuclei and synaptic ribbons in hair cells. Staining with an anti-FMRP antibody confirmed the presence of FMRP in the hair cells (inner and outer hair cells) of both young and adult WT mice (Fig. 1).

**Fig. 1.**
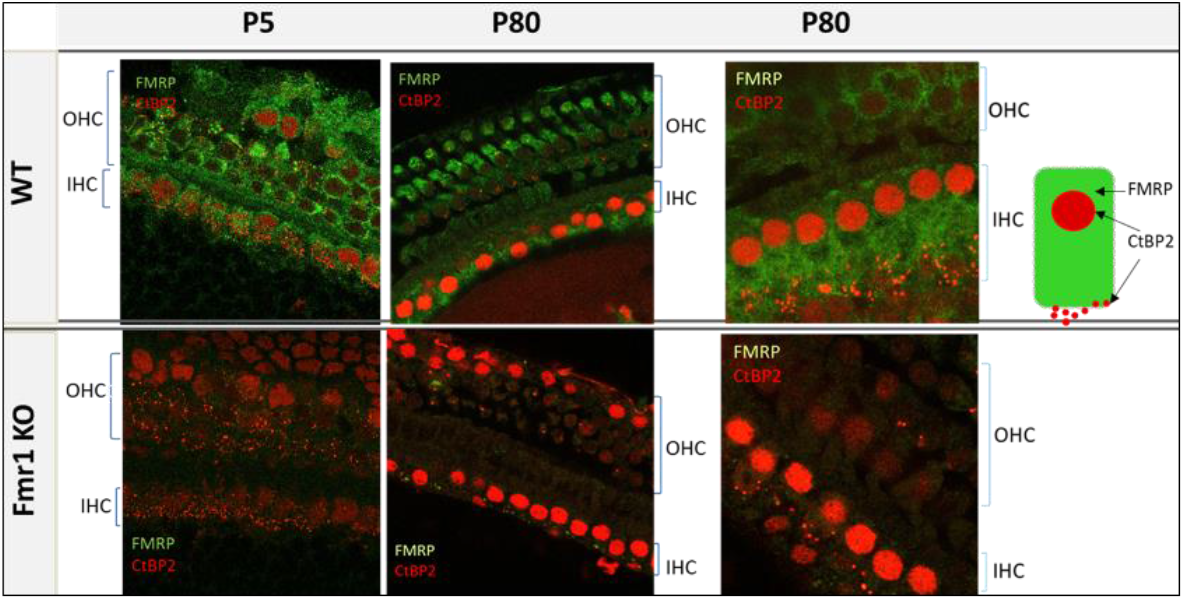
Representative images of () of the organ of Corti’s from FVB mice. IHC - inner hair cells, OHC - outer hair cells, P5 and P80, WT - upper panel, *Fmr1 KO* lower panel. Immunofluorescence staining using anti-CtBP2 (red) and anti-FMRP antibody (g reen), 63 x objective.

### Evaluation of the morphology of hair cells in Fmr1 KO and WT mice using confocal microscopy

The structural details of hair cells in *Fmr1 KO* mice cochlea have not been explored, therefore we started our analysis by examining the structure of inner hair cells (IHC) using confocal microscopy. We conducted immunofluorescence staining using specific antibodies that mark IHC and their pre- and postsynaptic compartments. The anti-CtBP2 antibody was utilized to stain both the synaptic ribbons and nuclei in the IHCs, while anti-Bassoon was used for localization of the presynaptic active zone protein Bassoon, and anti-GluA2 for the postsynapse. Additionally, phalloidin was employed to visualize the stereocilia, and myosin VI to stain the cell body. To observe mitochondria within the hair cells, we used the anti-Tom20 antibody (Fig. 2).

**Fig. 2.**
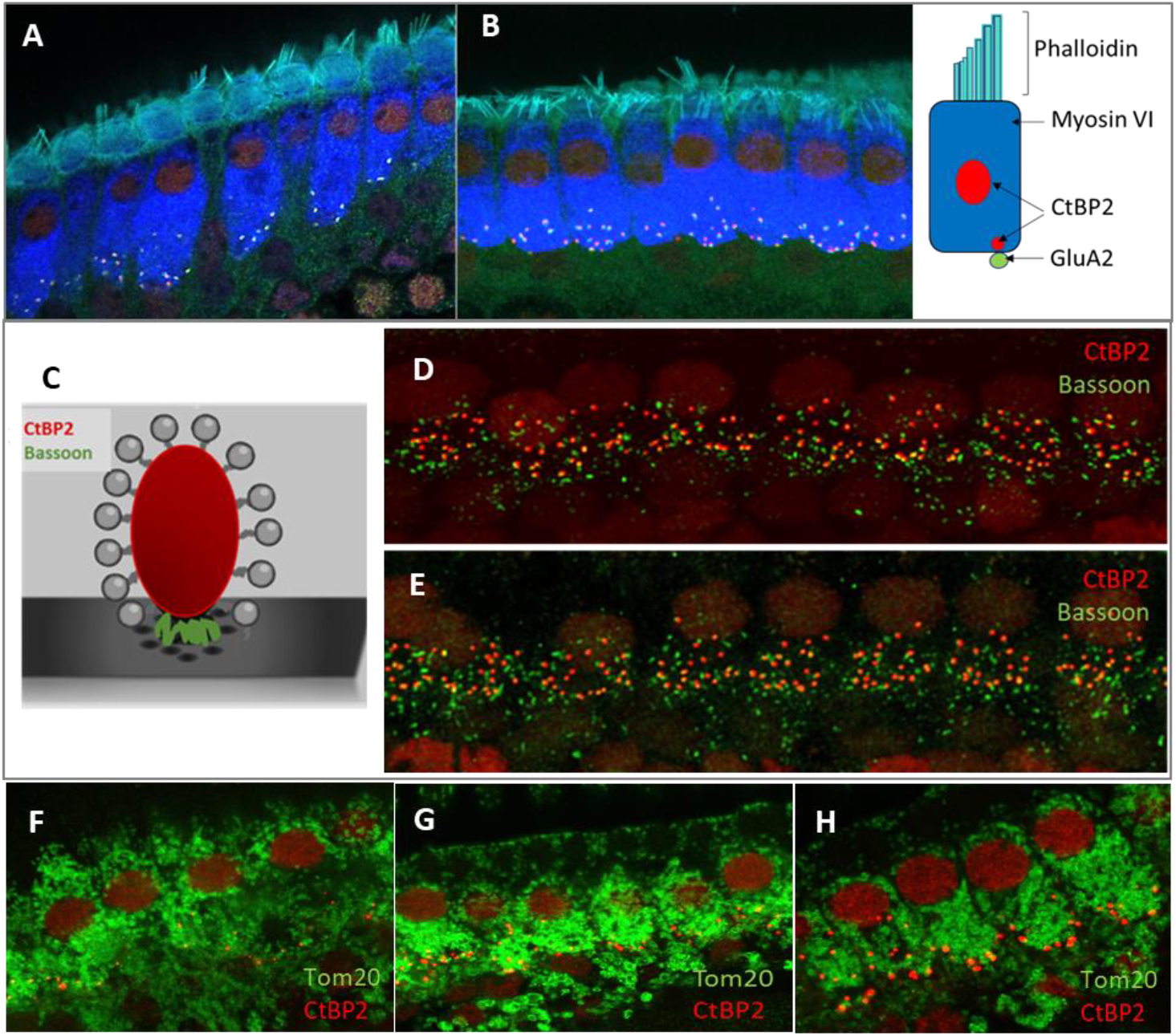
Analysis of the morphology of hair cells in adult *FMR1 KO* mice. Representative images of inner hair cells (IHC) of the organ of Corti’s from WT **(A)** and Fmr1FVB **(B)** mice. Schematic illustrations of ribbon synaps, a presynaptic markers, CtBP2(red) and Bassoon (green), placed between ribeye and the presynaptic membrane **(C)**. Immunostaining with anti-CtBP2 antibody (red) and anti-Bassoon antibody (green) in WT **(D)** and *Fmr1 KO* **(E)** mice. **(F, G, H)** Representative images of Tom20 as a marker for mitochondria in hair cells WT **(F)** and *Fmr1 KO* **(G)** mice, immunostaining with anti-CtBP2 antibody (red) and anti-Tom20 (FL-145) antibody (green). Enlargement of inner hair cells in *Fmr1 KO* mouse **(H)**, 63 x objective.

The confocal microscopy images revealed the typical morphology of the IHCs in the cochleae of *Fmr1 KO* mice. The stereocilia were situated on the apical part of the cell body, anchored to the cuticular plate (Fig. 2A, B), with the nuclei located in the apical half of the cell. Mitochondria were distributed throughout the cell body (Fig. 2F, G, H). The basal part of the IHCs housed multiple synaptic ribbons. Our findings show that the synaptic ribbons in IHCs from *Fmr1 KO* mice are positioned opposite the AMPA glutamate receptors (indicated by GluA2 staining) on the postsynaptic side, within the afferent nerve fibers (Fig. 2A, B and Fig. 8A, B, C).

Bassoon plays a crucial role in anchoring ribbons close to the presynaptic membrane. To study if its level is not changed in *Fmr1 KO* IHC we carried out immunostaining using anti-Bassoon and anti-CtBP2 antibodies. We observed that numerous Bassoon immunosignals overlapped with the ribbons in the IHCs of both WT and *Fmr1 KO* mice. Additionally, there were distinct Bassoon signals that did not overlap, which likely originated from efferent nerve terminals (Fig. 2D, E).

### Quantitative analysis of ribbon synapses in hair cells of Corti’s organ in Fmr1 KO and WT mice shows their delayed maturation

The maturation of ribbon synapses in inner hair cells (IHCs) is a complex process that is tightly regulated during early postnatal development (Michanski et al., 2019). To investigate how ribbon synapses develop in *Fmr1* knockout (KO) IHCs compared to wild-type (WT) mice, we conducted confocal microscopy analyses to assess their volume across three developmental stages, from postnatal day 5 (P5) through adulthood (Fig.3). These stages encompass both pre- and post-hearing onset periods.

To image the presynaptic ribbons we performed immunostaining using an anti-CtBP2 antibody on tissue isolated from the apical turns of the cochlea at P5, P14, and in adult mice (Fig. 3 D, E, F). This approach enabled us to quantify the number of ribbon synapses per IHC and assess the distribution of their average size.

**Fig. 3.**
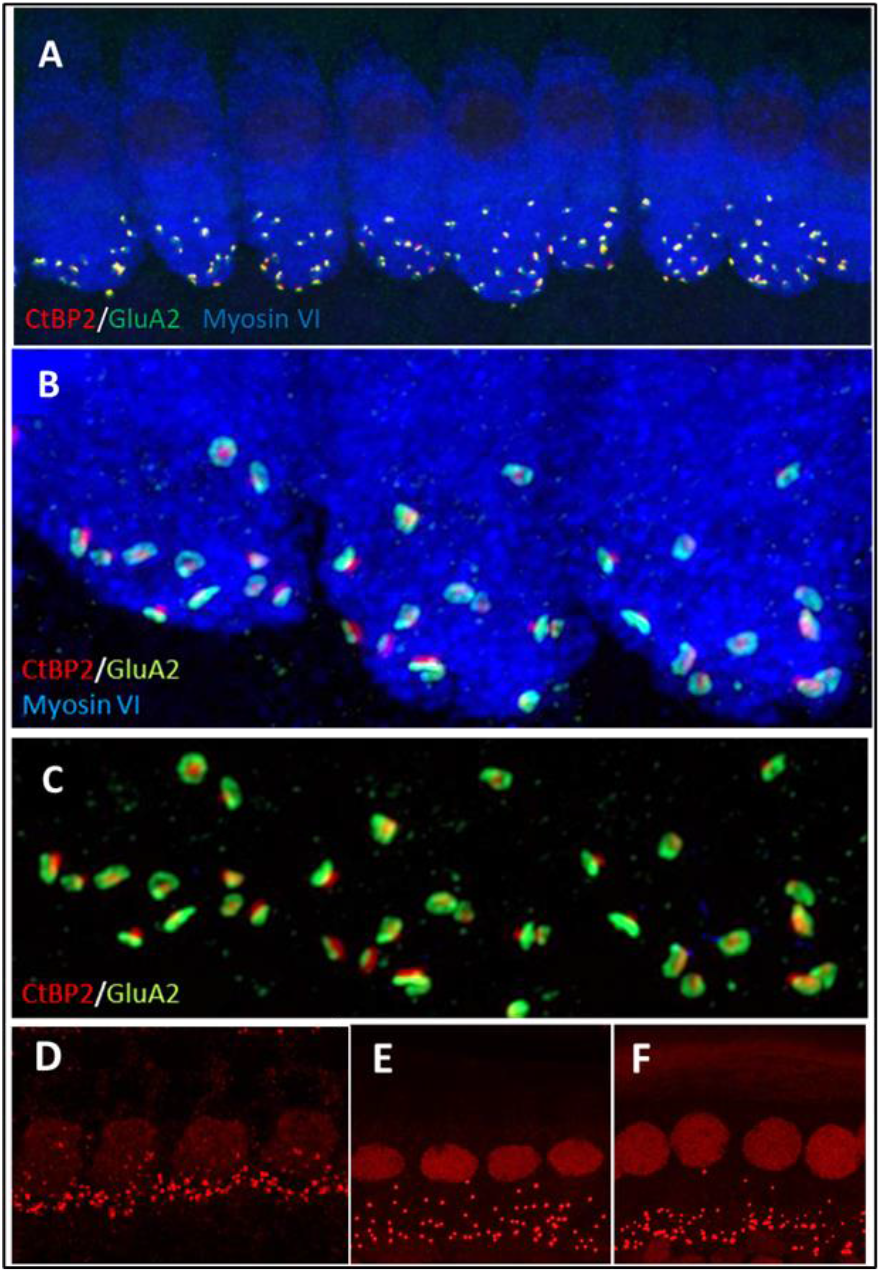
Immunofluorecent staining of IHCs and ribbon synapses in *Fmr1 KO* mice **(A)** Representative images of the IHC of the organ of Corti’s from P48 mice. White rectangular region is enlarged on (**B)** (63 x objective). The basal part of the cell containing multiple synaptic contacts, a presynaptic marker, CtBP2 (red dots), a postsynaptic marker, GluA2 (green dots). (**C)** Enlargement of the synaptic layer of IHC, synapses are visible as red and green dots. (**D, E, F)** Visualization of synaptic ribbons at P5, P14 and P48 hair cells in *Fmr1 KO* mice with anti-CtBP2 antibody. The CtBP2 protein is present in ribbon synapses and cell nuclei.

We observed that the number of CtBP2/RIBEYE puncta in IHCs significantly decreased from P5 to P14 (following the onset of hearing) and then stabilized. Although the number of synapses per IHC was comparable between wild-type and *Fmr1 KO* mice (Fig. 4 A, B, C), the average volume of fluorescently labeled presynaptic ribbons was notably smaller in P14 *Fmr1 KO* mice across three litters (Fig. 4 D, E, F).

**Fig. 4.**
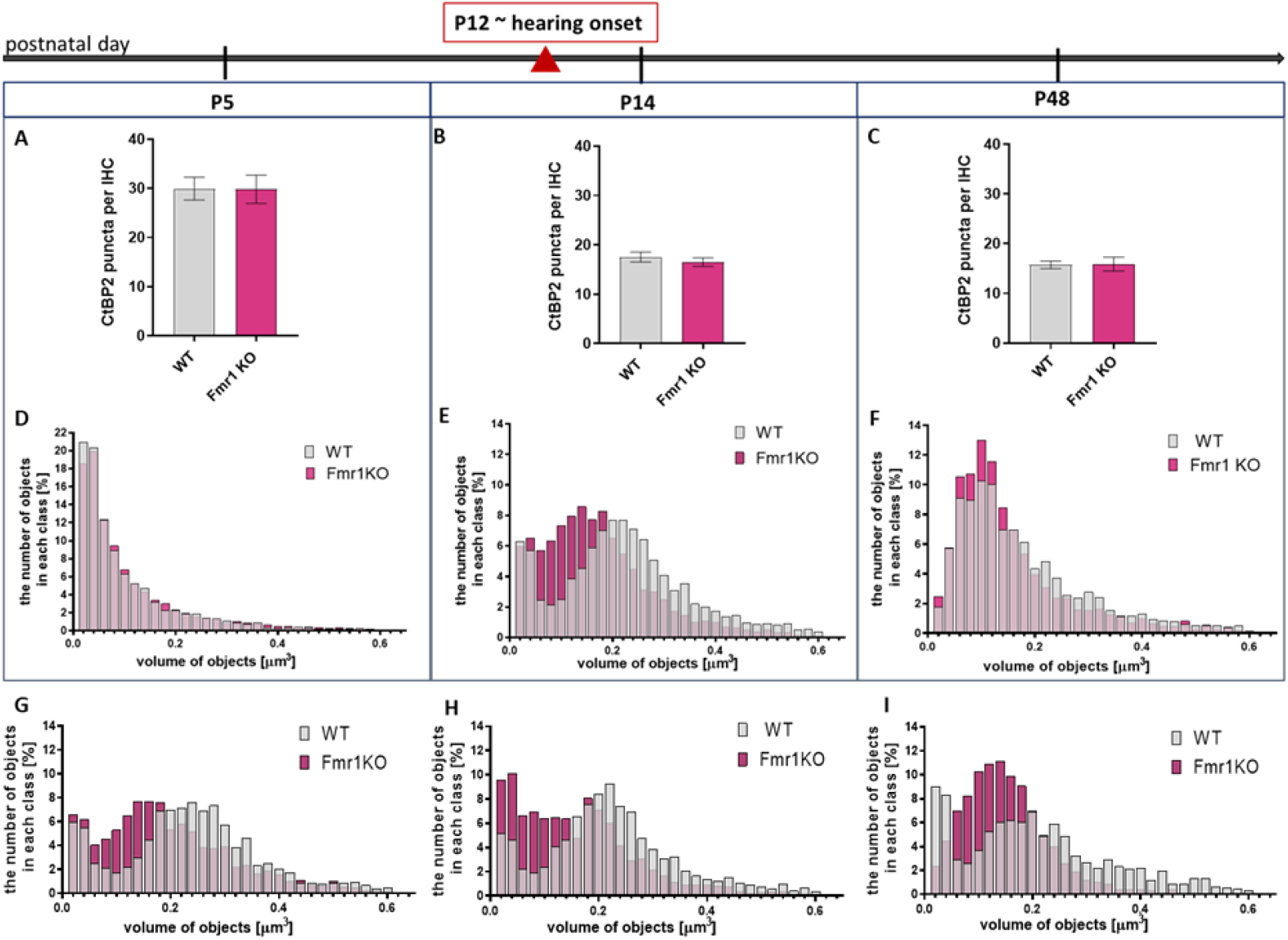
Changes in CtBP2/Ribeye in IHCs during development in apical turn of the cochlea Analysis of data based on anti-CtBP2 antibody staining against CtBP2/Ribeye protein (Fig.3) (**A, B, C)** Quantification of the number of CtBP2/Ribeye puncta per IHC at P5, P14, P48 WT and *Fmr1 KO* mice. Data is presented as mean, error bars indicate SEM, (t-test; P5 P-value = 0.9812; P14 P-value =0.4355, P48 P-value=0.9225) (**D, E, F)** The histograms presenting the distribution of synapses according to their volume in apical turn of the cochlea of P5, P14, P48 mice. Y axis –number of objects in each size class [%], X axis - volume of objects [μm3]. Number of animals: P5: N = 3 WT N = 5 *Fmr1 KO*, P14: N = 6 WT, N = 8 *Fmr1 KO*, P48: N = 4 WT, N = 4 *Fmr1 KO*. Data is presented as a relative frequency distribution histogram; P14 P-value <0.0001; P48 P-value <0.0001; Kolmogorov-Smirnov test. (**G, H, I)** The individual histograms for three independent litters of P14 mice (presented as combined data on panel E).

Histograms showing the distribution of average synapse sizes (categorized into arbitrarily defined size classes) revealed a higher prevalence of smaller ribbon synapses in the P14 *Fmr1 KO* (0.04–0.18 μm^3^), compared to a predominance of larger ribbons in wild-type litermates (0.2–0.6 μm^3^) (Fig. 4 E). Although this difference in ribbon volume was less pronounced in adult mice, smaller synapses remained more prevalent in *Fmr1 KO* mice at P48 (0.06–0.14 μm^3^) relative to wild-type controls (0.16–0.44 μm^3^) (Fig. 4 F).

Histograms representing the distribution of average synapse volume at P5 indicated that ribbons were comparable between wild-type and *Fmr1 KO* animals at early developmental stage before hearing onset (Fig. 4D).

### Inner hair cells ultrastructure in Fmr1 KO and WT mice using the transmission electron microscopy and the serial block face scanning electron microscopy

The ultrastructure of hair cells in *Fmr1 KO* mice has not been studied before. Therefore, to investigate the fine morphology of these cells, we used two complementary methods based on electron microscopy - the transmission electron microscopy (TEM) and serial block face scanning electron microscopy (SBF-SEM).

First, we evaluated the overall morphology of the organ of Corti in *Fmr1 KO* mice, as shown in the representative electron micrograph (Fig. 5A), and found no abnormalities in its organization. The sensory cells - inner hair cells (IHCs) and outer hair cells (OHCs) - were properly arranged along the basilar membrane, with the tectorial membrane positioned above them. Both IHCs and OHCs exhibited hair bundles, known as stereocilia.

**Fig. 5.**
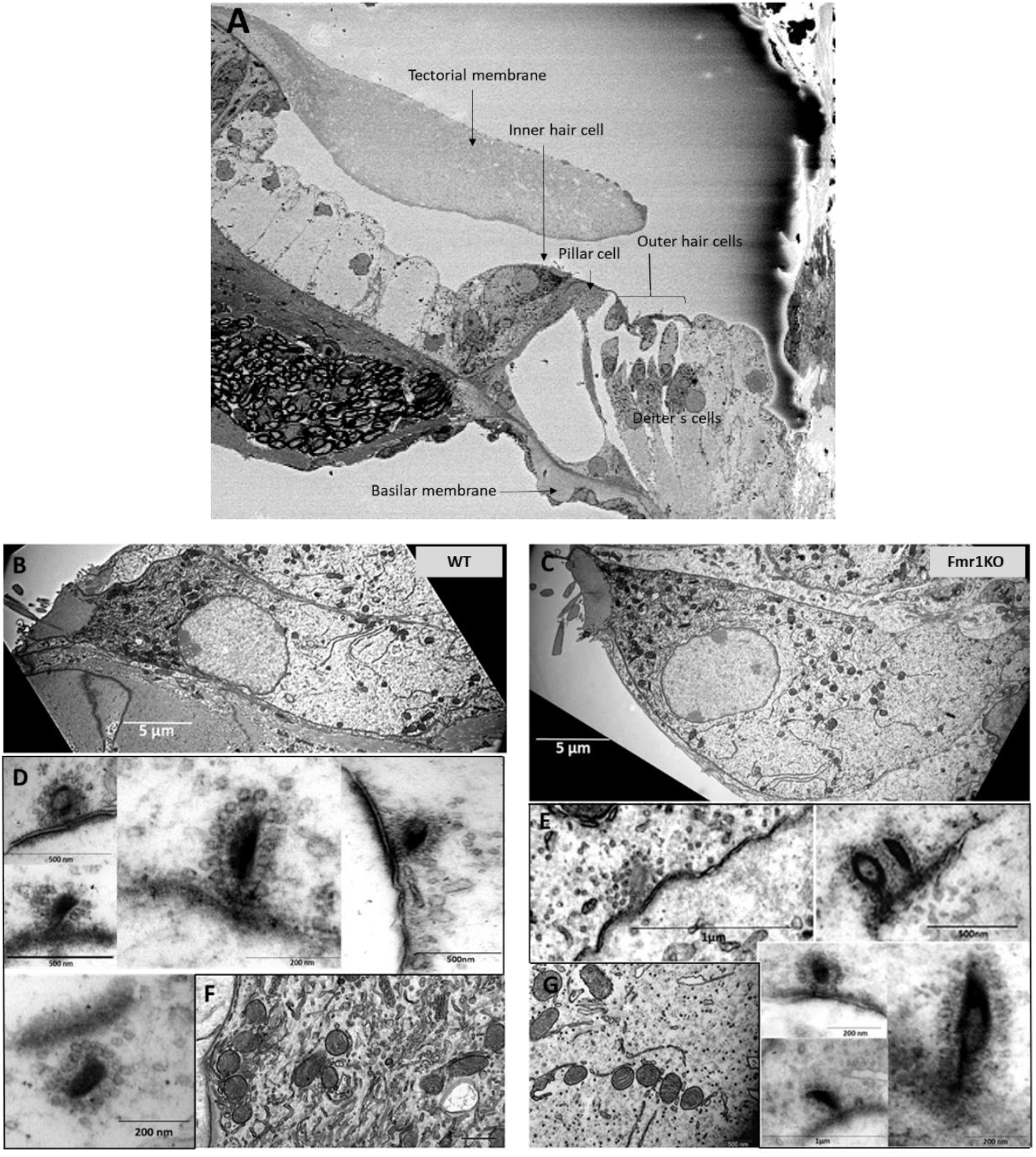
**(A)** Electron micrograph of the organ of Corti in *Fmr1 KO* mouse. **(B, C)** Ultrastructural analysis of cochlear inner hair cells. Representative electron micrographs of the WT (B) and *Fmr1 KO* inner hair cells **(C)** and their respective ribbon synapses surrounded by synaptic vehicles (synaptic vesicles are linked via filaments to the ribbon **D, E**) and mitochondria **(F, G)**.

Next, we investigated the ultrastructure of *Fmr1 KO* inner hair cells using transmission electron microscopy. Figure 4 presents representative images of IHCs from both *Fmr1 KO* and wild-type (WT) cochleae. The overall morphology of *Fmr1 KO* IHCs appeared comparable to that of WT cells, with well-defined stereocilia (Fig. 5 A, D). The shape of the *Fmr1 KO* IHCs, as well as the distribution of mitochondria and the nucleus, was similar to that observed in WT IHCs.

We observed typical postsynaptic densities and synaptic structures, including presynaptic dense bodies (ribbons) coated with synaptic vesicles. In some synapses, the ribbon was linked to the postsynaptic membrane via filamentous structures. Mitochondria were located beneath the cuticular plate and surrounding the nucleus, and were often associated with strands of intracellular membrane. The overall morphology of IHCs mitochondria appeared normal.

While transmission electron microscopy (TEM) provides high-resolution images of inner hair cells and ribbon synapses, it is limited in capturing the overall shape and morphology of entire cells. To address this question, we performed three-dimensional reconstructions of whole IHCs from both wild-type (WT) and *Fmr1 KO* mice using serial block-face scanning electron microscopy (SBF-SEM).

Based on serial image reconstructions, we extracted specific morphological parameters, such as the mean IHC surface area, which showed no significant differences between the two genotypes (Fig. 5B). We further reconstructed key cellular components, including the nucleus, mitochondria, synaptic ribbons, and nerve endings contacting individual IHCs. The mitochondrial distribution appeared uniform, and the localization of the nucleus was similar between *Fmr1 KO* and WT mice.

Typically, inner hair cells exhibit a distinct flask-like shape, characterized by a constriction in the neck region. However, in the *Fmr1 KO* mouse, the IHC displayed a more fusiform body shape compared to the wild-type control (Fig. 5). The nucleus was positioned in the apical half of the cell, while mitochondria were primarily concentrated beneath the cuticular plate and surrounding the nucleus.

Afferent terminals were identified by the presence of a postsynaptic density on the afferent ending, accompanied by a corresponding synaptic ribbon (Fig. 5, lower pannel). We analyzed the spatial distribution of synaptic structures surrounding the IHCs and performed three-dimensional reconstructions of ribbon synapses. These reconstructions did not reveal any notable dysmorphologies in *Fmr1 KO* mice compared to wild-type controls (Fig. 6).

**Fig. 6.**
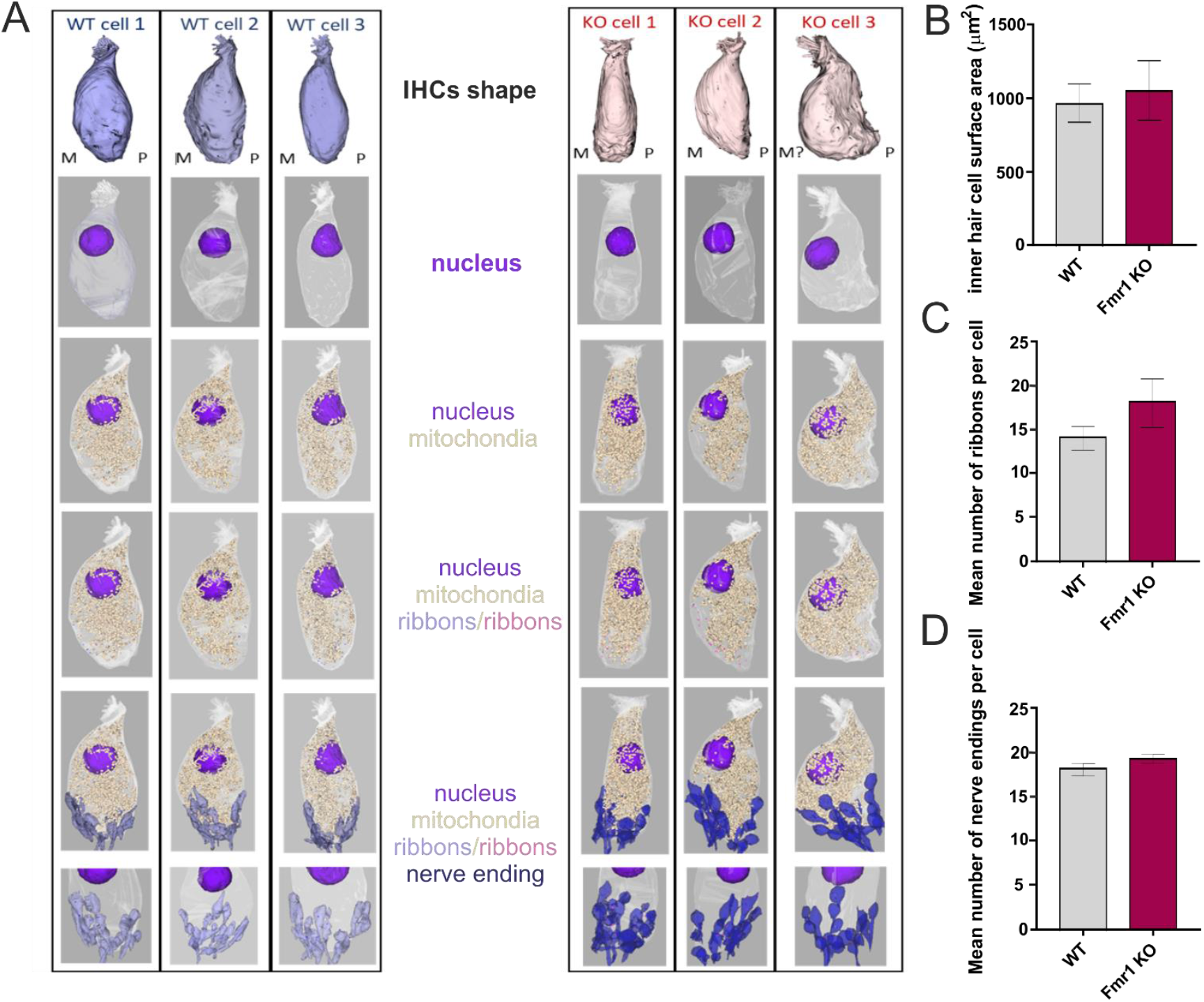
Detailed ultrastructural analysis of inner hair cells from adult WT (**A**, left pannel) and *Fmr1 KO* mice (**A**, right pannel). Three-dimensional reconstructions of IHCs shape, nucleus, mitochondria, ribbons and nerve endings were obtained by serial image reconstruction from SBF-SEM. **(B)** inner hair cell surface area WT vs *Fmr1 KO*, **(C)** mean number of ribbons per cell WT vs *Fmr1 KO*, **(D)** mean number of nerve endings pre cell WT vs *Fmr1 KO*. Data is presented as mean, error bars indicate SEM, Mann-Whitney nonparametric test P-value>0.05.

Quantitative analyses of both the number of ribbon synapses per cell (Fig. 5C) and the number of afferent nerve endings contacting each IHC (Fig. 5D) showed again no significant differences between WT and KO mice, indicating that the overall ultrastructure of IHCs in the inner ear remains intact in the absence of FMRP.

The quality of the SBF-SEM electron microscopy images was sufficiently good to enable detailed reconstruction of ribbon synapses for both mouse genotypes. For each genotype, we successfully reconstructed six synapses (Fig. 7). We measured both the surface area of each synapse and the number of synaptic vesicles associated with it (Fig. 7 B, C). These analyses revealed no statistically significant differences between *Fmr1 KO* and wild-type mice.

**Fig. 7.**
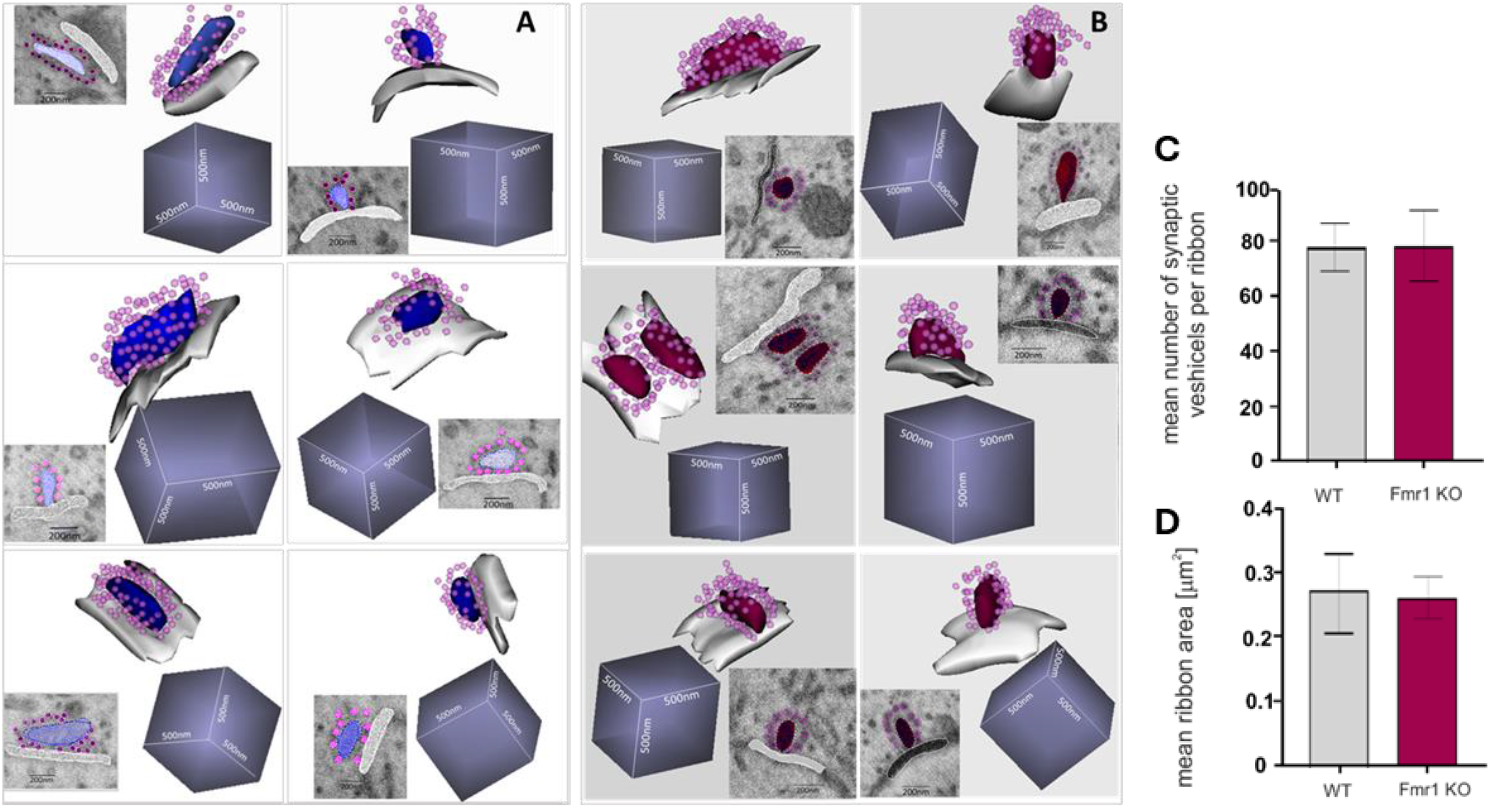
Ultrastructure of reconstructed ribbon synapses from WT **(A)** and *Fmr1 KO* **(B)** organ of Corti revealed by SBF-SEM. Mean number of synaptic vesicles per ribbon **(C)** and mean ribbon area **(D)** (n=6 synapses/genotype).

## Discussion

This study provides new insights into the morphological and ultrastructural features of *Fmr1 KO* IHCs ribbon synapses, the first connection on the path that conveys the auditory information from the sensory cells to the brain. Analysis at early postnatal stages (P5, P14) showed delayed developmental maturation of IHC ribbon synapses in Fmr1 KO mice. Interestingly the ultrastructure of IHCs and their ribbons studied by electron microscopy in the adult mice have shown no specific dysmorphologies in Fmr1 KO mice. These findings suggest that delayed maturation of auditory hair cell synapses in the absence of FMRP may contribute to atypical auditory circuit development driven by early sensory experience.

Altered synaptic structure and function is a major hallmark of FXS neurons; however, these changes have been studied primarily in brain synapses (Bagni and Zukin, 2019). Patients with FXS have been shown to experience difficulties processing auditory information in the cortical regions of the brain. Additionally, *Fmr1* knockout (*Fmr1 KO*) mice exhibit increased sensitivity to sound. Electrophysiological studies on both *Fmr1 KO* mice and FXS patients suggest that the impairment may occur at an early stage of auditory information processing, potentially beginning at the level of spiral ganglion neurons (Rotschafer and Cramer, 2017, Arinami et al., 1988).

ABR studies on *Fmr1 KO* mice revealed altered responses compared to wild-type mice, with changes evident from the first wave peak, linked to spiral ganglion neuron activity (Rattay and Danner, 2014). The peak amplitude was reduced, and the sound threshold needed to evoke a response was elevated in *Fmr1 KO* mice (Rotschafer et al., 2015), indicating impaired hearing. In humans with FXS, increased N1 and P2 ERP amplitudes suggest heightened responses from auditory and associative cortices (Van der Molen et al., 2012a). Moreover, the N1 amplitude does not decrease with repeated tones, indicating reduced adaptation to auditory stimuli (Castren et al., 2003). ERP studies consistently report elevated N1 amplitudes and reduced habituation to repeated sounds in FXS (Rojas et al., 2001, Castren et al., 2003, Van der Molen et al., 2012a, Van der Molen et al., 2012b, Schneider et al., 2013, Ethridge et al., 2016). MEG studies also show enhanced auditory responses in FXS (Rojas et al., 2001).

This led us to investigate synaptic connections in the inner hair cells of *Fmr1 KO* mice. We started by confirming FMRP expression in IHCs of Corti. In the brain, FMRP is expressed at high levels in auditory neurons at both cortical and subcortical levels (Beebe et al., 2014, Zorio et al., 2017, Yu et al., 2021). It was also shown to be present in hair cells and supporting cells of mice, rats, gerbils, and chickens, with a particularly high level in the immature hair cells during the prehearing period (Wang et al., 2023). We confirmed that FMRP is expressed in both inner and outer hair cells of WT mice, while *Fmr1 KO* mice lack this protein.

Since FMRP plays a critical role in synaptic development and plasticity, its absence may contribute to synaptic alterations in *Fmr1 KO* IHCs. We measured the number and volume of ribbon synapses in the IHCs of *Fmr1 KO* mice at the age of P5, P14 and P48. Before the onset of hearing, ribbon synapses in inner hair cells undergo significant molecular assembly and structural and functional maturation. During this process, synaptic contacts shift from multiple small active zones (AZs) to a single, larger one. This maturation involves the fusion of ribbon precursors with membrane-anchored ribbons, which can also merge with each other. These fusion events are most commonly observed around postnatal day 12 (P12), aligning with the onset of hearing in mice (Michanski et al., 2019).

We found that the average size of synaptic ribbons was significantly smaller in *Fmr1 KO* mice at P14 and remained slightly reduced in adulthood. That may reflect delayed maturation of ribbon synapses in the IHCs of *Fmr1 KO* mice. The total number of synapses per IHC was comparable between genotypes, but decreased with age as expected (Michanski et al., 2019). This suggests that FMRP is involved in the postnatal maturation of ribbon synapses, and its lack in Fmar1 KO mice is potentially affecting synaptic function and auditory processing. A higher proportion of smaller ribbons in *Fmr1 KO* mice indicates an alteration in synaptic developmental maturation, as ribbon size is linked to synaptic activity and neurotransmitter release efficiency (Mehta et al., 2013). The presence of GluA2-positive AMPA receptors at the postsynaptic side suggests that glutamatergic transmission is preserved; however, the observed differences in ribbon size in the critical age of development may contribute to abnormal development of circuits induced by auditory experience.

The overall morphology of IHCs did not differ between the genotypes. The localization of Bassoon, a key presynaptic scaffolding protein, overlapped with synaptic ribbons in both WT and *Fmr1 KO* mice, suggesting that FMRP deletion does not affect the anchoring of ribbons to the presynaptic membrane. However, distinct Bassoon-positive signals that did not co-localize with CtBP2-positive ribbons were detected, likely originating from efferent nerve terminals. This raises the possibility that alterations in efferent synaptic input may accompany the observed changes in ribbon synapse maturation.

Ultrastructural analyses using transmission electron microscopy (TEM) and serial block-face scanning electron microscopy (SBF-SEM) further revealed that the overall morphology of IHCs and the organ of Corti remained intact in *Fmr1 KO* mice. The spatial organization of stereocilia, mitochondria, and nuclei appeared similar between WT and *Fmr1 KO* IHCs, indicating that gross cellular morphology is preserved despite the absence of FMRP. In our previous study we detected abnormalities in the morphology of the mitochondria in *Fmr1 KO* brain (Kuzniewska B, 2019). Apparently in the IHCs of *Fmr1 KO* mice organ of Corti the mitochondria displayed perfectly normal shape and size and looked very similar to WT.

Three-dimensional reconstructions of ribbon synapses provided additional confirmation that the synaptic architecture of IHCs in *Fmr1 KO* mice remains largely unaltered. Synaptic ribbons were positioned opposite postsynaptic densities, and no major dysmorphologies were observed. Nevertheless, given the differences in ribbon size distribution, it is possible that synaptic function is subtly impaired, potentially contributing to auditory deficits commonly associated with fragile X syndrome.

Taken together, our findings indicate that *Fmr1 KO* mice exhibit a delay in ribbon synapse maturation without major structural abnormalities in IHCs. This suggests that FMRP is not essential for the initial formation of ribbon synapses but may play a crucial role in their postnatal development and refinement. Future studies should focus on functional assessments, such as electrophysiological recordings, to determine whether the observed morphological changes translate into synaptic transmission deficits. Understanding the impact of FMRP loss on auditory processing at a cellular level could provide valuable insights into the auditory impairments seen in fragile X syndrome and related neurodevelopmental disorders.

## Acknowledgments

This work was supported by NCN grant 2017/27/B/NZ4/01169 for MD.

We would like to thank Dr Piotr Kaźmierczak and Dr Maciej Winiarski for methodological support in this project.

## Author contributions

Conceptualization and supervision: MD.; Validation: All authors participated in the interpretation of the data; Formal analysis: MC; Investigation: MC, AS; Writing - original draft: MD, MC, AS; All authors participated in the manuscript review and editing; Funding acquisition: M.D.

